# L1CAM-CAR T cells with enhanced potency overcome low-density antigen expression in rhabdomyosarcoma

**DOI:** 10.64898/2026.02.27.707949

**Authors:** Caroline Piccand, Corentin Gauthier, Sara G. Danielli, Rhoikos Furtwängler, Jochen Rössler, Andrea Timpanaro, Michele Bernasconi

**Affiliations:** Department of Pediatric Hematology and Oncology, Inselspital, Bern University Hospital, 3010 Bern, Switzerland; Translational Cancer Research, Department for BioMedical Research (DBMR), University of Bern, 3008 Bern, Switzerland; Graduate School for Cellular and Biomedical Sciences, University of Bern, 3012 Bern, Switzerland. Ben Towne Center for Childhood Cancer and Blood Disorders Research, Seattle Children’s Research Institute, Seattle, 98101 WA, USA; Department of Oncology and Children’s Research Center, University Children’s Hospital of Zurich, Zürich 8032, Switzerland

**Author notes:** Corresponding author (M.B.). (C.P.); (C.G); (R.F.); (J.R.); (A.T.), (S.D).

**Keywords:** Chimeric antigen receptor, L1CAM, CD171, Rhabdomyosarcoma, Immunotherapy, B7-H3, CD276

## Abstract

Rhabdomyosarcoma (RMS), the most common pediatric soft tissue sarcoma, shows dismal survival in relapsed or metastatic alveolar disease. Chimeric antigen receptor (CAR) T cells are promising but limited by scarce tumor-selective antigens and suboptimal efficacy at low antigen density.

We investigated L1 cell adhesion molecule (L1CAM) as a therapeutic target by profiling its expression by flow cytometry, immunoblotting, and immunohistochemistry in cell lines, patient-derived xenografts, and healthy tissues. Using the scFv derived from the CE7 antibody, we engineered L1CAM-CARs with distinct hinge and costimulatory domains and tested them *in vitro* and in orthotopic RMS mouse models against clinically tested CE7- and B7-H3-CARs. L1CAM was consistently expressed at moderate levels in RMS, especially alveolar subtypes, but very weakly expressed in healthy tissues. Flow cytometry revealed a moderate density typically limiting CAR activity. Among constructs, L1CAM.III (CE7-CAR with long hinge and CD28 domain) showed the strongest cytotoxicity and IFN-γ release. *In vivo*, L1CAM.III-CAR T cells regressed tumors, prolonged survival, and persisted in orthotopic RMS models, showing greater efficacy in alveolar RMS and no off-tumor activity.

These findings establish L1CAM as a rational RMS therapeutic target. Optimized L1CAM.III-CAR T cells overcome moderate antigen density, achieving potent and persistent antitumor activity comparable to B7-H3-CARs but with improved safety. This work supports CAR optimization for clinical translation to broaden pediatric sarcoma immunotherapy.

## 1. Introduction

Rhabdomyosarcoma (RMS) is the most common soft tissue sarcoma, accounting for nearly half of malignant sarcomas in children and adolescents.^1^ RMS arises from primitive skeletal muscle progenitors and is histologically classified into embryonal (eRMS) and alveolar (aRMS) subtypes. More recently, molecular stratification based on PAX3/7::FOXO1 gene fusions distinguishes fusion-negative (FN) from fusion-positive (FP) tumors.^2,3^ FP-RMS, typically of the alveolar subtype, displays particularly aggressive clinical behavior and carries a poor prognosis, whereas FN-RMS, including most embryonal cases, is associated with more favorable outcomes. Despite multimodal therapy, survival for relapsed or metastatic FP-RMS remains below 30%,^4^ underscoring the urgent need for improved treatments.

Due to the remarkable results in the treatment of hematologic cancers^5,6^ and their translation into pediatric solid tumors - including RMS^7–11^ - our focus has shifted to immunotherapies, especially chimeric antigen receptor (CAR) T cell therapy. Preclinical studies in RMS have demonstrated encouraging antitumor activity of CAR T cells targeting specific antigens, such as HER2,^7^ B7-H3,^9,11,12^ or FGFR4^8–11^. In the clinical setting, HER2-targeting CAR T cells were evaluated in pediatric and young adult sarcomas (NCT00902044), demonstrating feasibility and acceptable safety.^13^ Among three RMS patients treated, two experienced disease progression and subsequently died, whereas one patient achieved a complete response and has remained tumor-free for 71 months upon CAR T cell treatment.^7^ These findings are very encouraging, but highlight the need for further optimization of CAR T cell therapies in RMS. To this end, a phase I trial (NCT04995003) is currently evaluating HER2-CAR T cells in combination with immune checkpoint inhibitors (pembrolizumab or nivolumab) in patients with advanced sarcomas, including RMS, aiming to enhance CAR T cell expansion, persistence, and therapeutic efficacy.

Tumor-associated antigens are often expressed in healthy tissues, raising the risk of on-target/off-tumor toxicity.^14^ By contrast, tumor-restricted antigens tend to be expressed at lower density or heterogeneously, thus limiting CAR T cell activation and tumor killing.^9,11^ Designing CAR T cells that achieve potent, yet selective antitumor activity is therefore critical to develop safe and effective immunotherapy options for RMS patients.

Previously, we identified RMS-associated targets that are upregulated compared to healthy tissues. Among these, the L1 cell adhesion molecule (L1CAM) showed particularly high expression and was correlated with lower survival in the aggressive alveolar RMS.^15^ L1CAM is a transmembrane glycoprotein involved in neural adhesion, migration, and signal transduction.^16^ Beyond its critical role in nervous system development, L1CAM expression has been significantly associated with worse survival, tumor progression, metastasis, and therapy resistance across multiple malignancies^17^, including neuroblastoma,^18^ ovarian cancer,^19^ retinoblastoma,^20^ and melanoma.^21^

Preclinically, L1CAM-CAR T cells have shown activity in neuroblastoma,^18^ retinoblastoma,^20^ and ovarian cancer models.^22^ Importantly, L1CAM was already validated as a target in a phase I clinical trial for neuroblastoma (NCT02311621), which demonstrated tolerability in patients and CAR T cell trafficking to tumor sites.^23^ While no objective responses were achieved, the trial confirmed the feasibility of targeting L1CAM in children and established it as a clinically validated target, underscoring the need to refine CAR designs to enhance its full therapeutic efficacy. To date, L1CAM-targeted CAR T cells have not been tested in RMS.

Both the hinge region and costimulatory domain (CSD) are key determinants of CAR T cell performance. The hinge can influence epitope accessibility and immune synapse formation,^24^ whereas the CSDs support persistence and activation thresholds.^25^ Notably, *in vitro* studies have shown that CARs with CD28-derived CSD deliver stronger early activation than those with 4-1BB, which may be particularly advantageous under conditions of low antigen density.^26^

In this study, we investigate L1CAM as a therapeutic target in RMS by assessing its cell-surface expression across multiple RMS models and its distribution in healthy tissues. We further evaluate optimized L1CAM-CAR T cells, designed to maximize antitumor activity while minimizing off-tumor toxicity, *in vitro* and in orthotopic RMS models.

## 2. Material and methods

### Cell culture

Fusion-negative (RD, Rh18, Rh36, TTC442, RUCH-3) and fusion-positive (Rh4, Rh5, Rh28, Rh30, JR, RMS) RMS cell lines (kindly provided by Dr. Marco Wachtel, University Children’s Hospital Zurich) were cultured in DMEM supplemented with 10% FBS, 2 mM L-glutamine, and 100 U/mL penicillin/streptomycin at 37 °C and 5% CO₂. Two aRMS patient-derived xenografts (IC-pPDX-104 and RMS-ZH003) were included and maintained as previously described. MRC-5 fibroblasts (ATCC) and immortalized human myoblasts (KM155C25Dist) were cultured in Skeletal Muscle Cell Growth Medium with SupplementMix. Firefly luciferase-expressing RMS cell lines (fLuc⁺) were maintained in RPMI-1640 with identical supplements.

### L1CAM knockout RMS cell lines

sgRNAs targeting exon 4 of L1CAM were designed using the GUIDES tool, as previously described.^9^ Two sgRNAs (5′-CACCGCCTGCTTATCCAGATCCCCG-3′ and 5′- AAACCGGGGATCTGGATAAGCAGGC-3′) were cloned into pLentiGuide-PURO,^27^ a gift from Feng Zhang (Addgene, Watertown, MA, #52963). The pLentiCRISPR-mCherry vector^28^, a gift from Beat Bornhauser (Addgene, #75161) was used to stably express Cas9 and mCherry in RD and Rh30 cells. Following lentiviral transduction (MOI 5), mCherry⁺ cells were FACS-sorted, transiently transfected with pLentiGuide-PURO, selected with puromycin (1 µg/mL) (InvivoGen, San Diego, CA) and expanded as single-cell clones. The targeted region was PCR-amplified, TOPO-cloned, and Sanger sequenced to confirm frameshift mutations.

### Truncated mouse L1CAM expressing RD and Rh30 cell lines

A truncated mouse L1CAM sequence (retaining the intracellular KRS motif) was PCR-amplified from MG50835 (Sino Biological Inc., Beijing, China) and ligated into pCDH-EF1 (Addgene) for lentivirus production. fLuc^+^ L1CAM knockout RD and Rh30 cells were transduced with the resulting lentivirus at a MOI of 5. 48 h later, cells were sorted for mouse L1CAM expression using PE- conjugated anti-mouse CD171 (L1CAM) antibody (1:100, Miltenyi Biotec, Bergisch Gladbach, Germany) to have a homogenous population.

### CAR constructs generation

The CE7 scFv, specific for L1CAM,^29^ L1CAM CT CAR plasmid (pj03196), was kindly provided by Prof. Michael C. Jensen (Seattle Children’s Research Institute, Seattle, WA) and used to replace the FMC63 scFv in the CD19 CAR backbone (pLV-EF1A-CD19.28H.28TM.28.3z, Addgene, #200678) via Esp3I digestion (Thermofisher) and ligation. All constructs included an N-terminal MycTag for standardized surface detection. Alternative hinges (CD28 or IgG4CH2CH3) and costimulatory domains (CD28 or 4-1BB) were PCR-amplified from existing templates^9^ and assembled into the CE7 CAR backbone using Gibson assembly (Thermofisher), generating a panel of CE7-based CARs. A B7-H3-targeting CAR was generated using the same scFv-swapping strategy with the CD276.MG scFv, kindly provided by Prof. Robbie Majzner (Stanford University, Stanford, CA).

Antares⁺ CAR constructs were generated by replacing the eGFP domain via BsiWI and ApaI digestion (Thermofisher), followed by Gibson assembly using a PCR-amplified P2A fragment and the Antares luciferase sequence (pNCS Antares, Addgene, #74279). Antares, a NanoLuc-based red-shifted reporter, enables sensitive in vivo imaging of CAR T cell dynamics. All plasmids were transformed into One Shot TOP10 E. coli (Thermofisher), purified using NucleoBond Xtra Midi kits (Macherey-Nagel), assessed by NanoDrop, and sequence-verified by Sanger sequencing (Microsynth).

### Lentivirus production

Lentiviral particles were produced by transient co-transfection of Lenti-X 293T cells (Takara Bio Inc., Kusatsu, Japan) on collagen type I rat tail-coated dishes (1:10 in sterile water, Sigma-Aldrich). 5×10^6^ cells were transfected the following day using the calcium phosphate method as previously described^9^. Supernatants were collected 48 h later, filtered (0.45 µm), and concentrated by overnight centrifugation at 4’000x*g* at 4 °C.

### CAR T cell manufacturing

CAR T cells were generated from donor PBMCs using our previously described protocol⁹, including CD3/CD28 activation, lentiviral transduction, and expansion in IL-2/IL-21-supplemented medium. Functional assays were performed with day-14 CAR T cells; for in vivo studies, either freshly manufactured (Rh30 model) or cryopreserved day-14 cells (RD model; viability >85%) were used. Day-14 products showed ∼50:50 CD4/CD8 ratios and minimal CD69 expression

### *In vitro* cytotoxicity assays

fLuc^+^ RMS cells were generated as previously described^9^. For bioluminescence-based killing assays, 5×10^3^ fLuc^+^ RMS target cells were seeded 24 h in advance in white 96-well plates suitable for luminescence detection (Thermofisher). 4 h before co-culture in white 96-well plates (Thermofisher). CAR T cells were added at E:T ratios of 1:8, 1:4, 1:1, and 2:1 in 200 µL RPMI-1640. After 48 h, 1.5 µg/well D-luciferin (VivoGlo^TM^ Luciferin, In Vivo Grade, Promega, Madison, WI) was added and bioluminescence measured on a GloMax luminometer (Promega). Data from three donors (technical triplicates) were analyzed using Excel and GraphPad Prism 10 and expressed as percent survival relative to target-only controls.

### Cytokine release assays

RMS target cells (75×10^3^/well were seeded in U-bottom 96-well plates and cocultured with CAR T cells at a 2:1 E:T ratio in 200 µL RPMI-1640. After 24 h, supernatants were collected, centrifuged, and stored at −80 °C. IFN-γ levels were quantified using a human IFN-γ ELISA kit (Invitrogen/Thermofisher) according to the manufacturer’s instructions. Data from three independent donors (technical triplicates) were analyzed in Microsoft Excel and plotted with GraphPad Prism 10.

### Western blotting

Cells were lysed in RIPA buffer with Halt™ protease inhibitors (Thermo Fisher), incubated on ice, and sonicated. Protein concentrations were measured using the Pierce BCA assay (Thermo Fisher), and 30 µg lysates were resolved on SurePAGE™ Bis-Tris 4-12% gels (GenScript). Membranes were probed overnight at 4 °C with anti-L1CAM (UJ127.11, 1:1,000; Thermo Fisher) or anti-α-tubulin (1:1,000; Cell Signaling), followed by HRP-conjugated secondary antibodies (1:10,000; Cell Signaling). Signals were detected with SuperSignal West Femto (Thermo Fisher) on a ChemiDoc MP system (Bio-Rad). Band intensities were quantified in Image Lab (Bio-Rad) and normalized to α-tubulin.

### Flow cytometry

Surface L1CAM expression was assessed after Accutase detachment (Sigma-Aldrich) by staining in FACS buffer (PBS + 2% BSA) with PE-anti-CD171/L1CAM REAfinity™ (1:100; Miltenyi) or rabbit anti-L1CAM CE7 (0.25 ng/µL; ABCD Antibodies) followed by Alexa Fluor™ 488 goat anti-rabbit IgG (1:1,000; Thermo Fisher). Data were acquired on a CytoFLEX cytometer (Beckman Coulter) and analyzed in FlowJo v10.8.1 (BD). Antigen density was quantified using QuantiBRITE™ PE beads (BD). CAR T cells were stained with BD Multitest CD3/CD4/CD8/CD45 (1:20; BD) and APC-anti-CD69 (1:100; BD) in FACS buffer.

### Immunohistochemistry

IHC was performed on FFPE sections as previously described,^9^ using citrate-based antigen retrieval and DAB detection. Slides were incubated overnight at 4 °C with anti-L1CAM (1:50; Sigma-Aldrich) or anti-CD3 (1:200; Abcam), followed by HRP-conjugated secondary antibodies (Agilent). Staining was developed with DAB, counterstained with hematoxylin, and imaged on a Leica DM6000 B microscope.

TMA staining intensity was quantified in QuPath v0.5.1 using H-scores, categorized as low (≤50), moderate (51-149), or high (≥150). Healthy tissue TMAs were analyzed in ImageJ using color deconvolution, DAB intensity binning (0-55 negative; 55-70 low; 70-110 medium; 110-255 high), and H-score calculation (%low×1 + %medium×2 + %high×3). Values were normalized to the highest-scoring sample (=100).

### Tissue microarrays

We built a tissue microarray of RMS PDXs by collecting or generating RMS PDXs as previously described.^30^ When reaching a size of 700-1’300 mm^3^, tumors were isolated from mice, mechanically minced into smaller pieces using scalpels, and fixed in ROTI Histofix 4% (Carl Roth, P087.3).

### *In vivo* orthotopic RMS models

Female NSG mice (4-6 weeks old; NOD.Cg-Prkdc^SCID^Il2rg^tm1Wjl^/SzJ; Charles River, Sulzfeld, Germany) were housed under L conditions at the University of Bern. All procedures were approved by the Canton Bern Animal Welfare Office (BE22/2022). After 2 weeks of acclimatization, mice were injected intramuscularly into the left gastrocnemius with 5×10⁵ fLuc⁺ RD or 2.5×10^5^ fLuc⁺ Rh30 cells in 50 µL PBS. Tumor burden was assessed on day 4 by bioluminescence imaging (IVIS Spectrum, PerkinElmer). Mice received 5×10^6^ CAR T cells intravenously on days 5 and 12. fLuc⁺ RMS cells were visualized using intraperitoneal VivoGlo™ Luciferin (Promega), and nanoLuc⁺ CAR T cells using Nano-Glo® In Vivo Substrate (Promega). Data were analyzed with Living Image® (PerkinElmer) and Microsoft Excel, and figures generated in GraphPad Prism 10.

### Statistical analysis

Statistical analyses were performed in GraphPad Prism 10. Killing and cytokine assays were evaluated by two-way ANOVA with Dunnett’s multiple comparison test. Statistical significance was reported as follows: *p* > 0.05 (ns), *p* ≤ 0.05 (*), *p* ≤ 0.01 (**), *p* ≤ 0.001 (***), and *p* ≤ 0.0001 (****).

## 3. Results

### L1CAM is expressed in multiple RMS cell lines and PDXs

Given the robust expression of L1CAM transcripts across multiple RMS cell lines and lower levels in healthy extracranial tissues, together with the consistent expression detected by IHC in 16/19 aRMS patients’ samples,^15^ we tested L1CAM protein expression in multiple RMS models. Western blot analyses revealed elevated L1CAM protein levels in most patient derived xenografts (PDXs) and cell lines, with notably high levels in FN-RMS Rh18, TTC442, RD, and Rh36. Overall, expression was higher in FN-RMS cell lines, compared to FP-RMS cell lines. In contrast, immortalized human myoblasts, MRC-5 fibroblasts, peripheral blood mononuclear cells (PBMCs), and T cells showed minimal signal for L1CAM (Figure 1A). Flow cytometry further demonstrated uniform L1CAM surface expression across RMS cell lines, and Quantibrite assessment estimated 2’000-20’000 L1CAM molecules per cell. As expected, control cells showed a minimal signal (Figure 1B).

**Figure 1.**
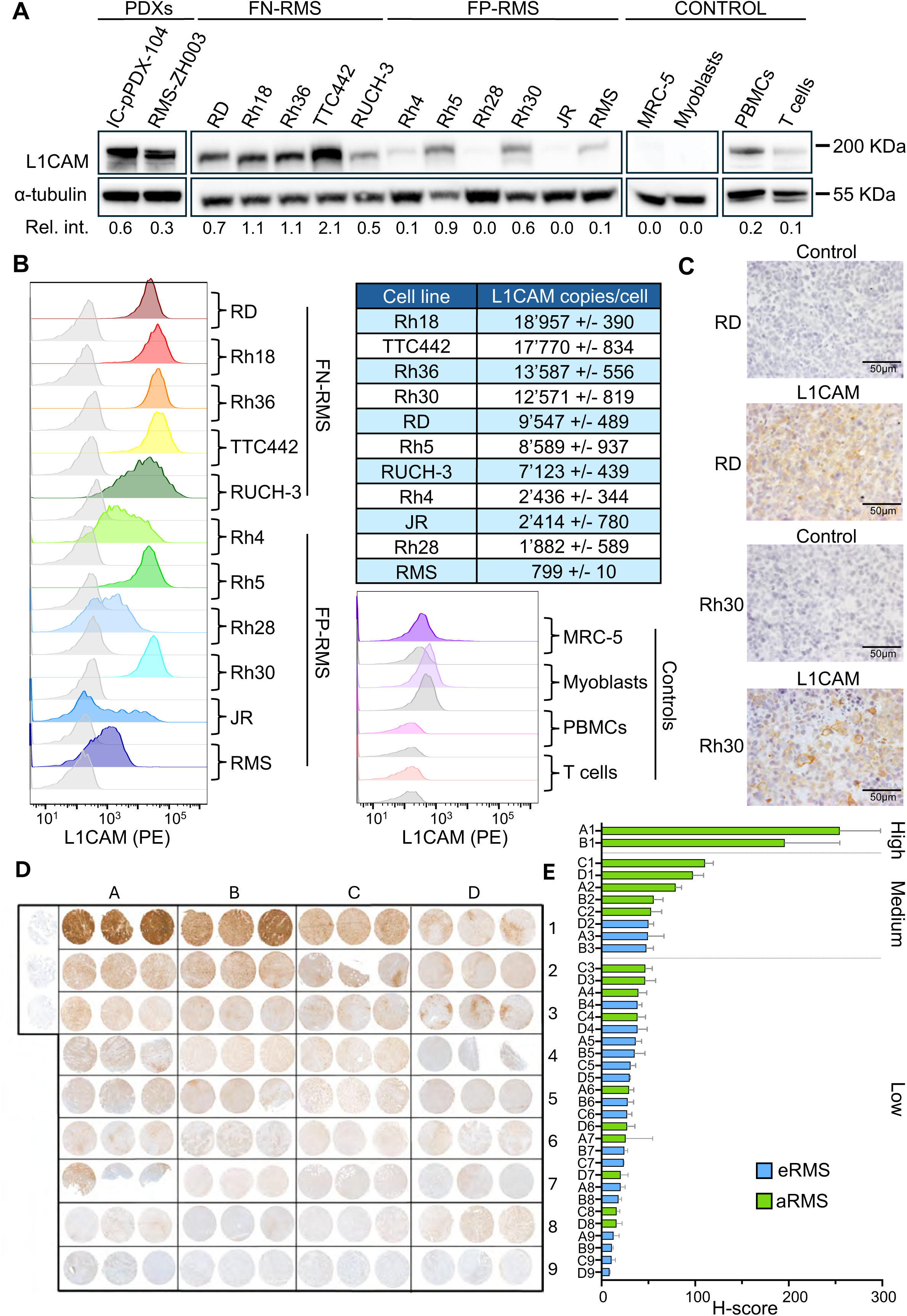
L1CAM is expressed in RMS models with heterogeneous levels across PDXs. **(A)** Western blot of L1CAM (∼200 kDa) in RMS PDXs and FN-/FP-RMS cell lines; *α*-tubulin served as loading control. Numbers below lanes indicate relative intensities normalized to α-tubulin. **(B)** Flow cytometry of FN- and FP-RMS cell lines and controls (gray isotype). Antigen density estimated with Quantibrite-PE calibration. Data representative of three independent experiments. **(C)** IHC of orthotopic xenografts (RD, Rh30) with L1CAM staining (brown) and hematoxylin counterstain (blue). Negative-control sections without primary antibody. Scale bars, 50 µm. **(D-E)** Tissue microarray of 34 primary RMS PDXs stained for L1CAM (brown), with triplicate tumor cores per case. The array included 36 cores from 34 xenografts; cores A9/C7 and A6/B2 were duplicates. (E) H-score quantification of L1CAM expression per case. Expression categorized as High (≥150), Medium (51-149), or Low (≤50). Subtypes color-coded (aRMS, green; eRMS, blue).

To validate our i*n vivo* models, we performed immunohistochemical (IHC) staining on orthotopic RMS xenografts derived from Rh30 and RD cells, the most representative for aRMS and eRMS subtypes, respectively. Rh30 xenografts showed stronger predominantly membranous staining with focal regions of high intensity, whereas RD xenografts displayed diffuse moderate membranous and cytoplasmic L1CAM staining (Figure 1C). These patterns confirm L1CAM expression in both RMS models, supporting their suitability for preclinical evaluation of L1CAM-targeted therapies.

To further assess the translational relevance of L1CAM in the context of RMS, we analyzed L1CAM expression in a tissue microarray (TMA) of 34 primary RMS patient-derived mouse xenografts (Figure 1D). L1CAM expression across RMS PDXs was heterogeneous, with strong staining detected in a minority of aRMS cases and moderate staining present in subsets of both aRMS and eRMS models. Specifically, 2/17 aRMS PDXs revealed high-grade staining (H-score ≥ 150), and 5/17 aRMS PDXs showed moderate staining (50 < H-score ≤ 149) (Figure 1E, green). Notably, 3/19 eRMS cases displayed moderate positivity (Figure 1E, blue).

Together, these findings demonstrate that L1CAM is expressed across RMS cell lines and in a subset of RMS PDXs, with higher frequencies in aRMS and detectable expression in some eRMS models.

### Generation and optimization of L1CAM-targeting CAR T cells

Several L1CAM-targeting antibodies have been described, including UJ127,^31^ L1-11A,^32^ and CE7^29^ - that recognize distinct extracellular L1CAM epitopes and could provide an antigen-binding domain for CAR design. Although L1CAM is broadly glycosylated, tumors can display a unique *N*-acetylglucosamine-containing epitope that is recognized by the CE7 antibody but is absent in healthy tissues.^33–35^ CE7 has translational relevance as it has been used to stain healthy and rhesus macaque tissue arrays, showing similar expression, and the safety of CE7-based L1CAM-CAR T cells has been validated in non-human primates.^18^ These CE7-based anti-L1CAM CAR T cells were then tested in a Phase I trial for pediatric neuroblastoma (NCT02311621).^23^ Building on this rationale, we examined CE7 antibody binding in RMS cell lines. Flow cytometry showed strong CE7 reactivity on the surface of Rh30 and RD cells, with no signal detectable in the CRISPR/Cas9-derived L1CAM knockout counterparts^36^ (Figure 2A). We therefore used the CE7-derived single-chain variable fragment (scFv) as the antigen-binding domain to generate L1CAM-directed CAR T cells, using CAR backbone that had previously demonstrated high efficacy against multiple RMS models.^9^ We replaced the B7-H3 antigen-binding domain with the CE7 scFv, yielding the construct CE7scFv.CD28H.CD28TM.CD28.CD3ζ (hereafter referred to as “L1CAM.III”). As a reference, we included the clinically validated L1CAM-CAR (CE7scFv.IgG4CH2CH3.CD28TM.4-1BB.CD3ζ, hereafter “L1CAM.CT”) tested in the Phase I neuroblastoma trial.^23^

**Figure 2.**
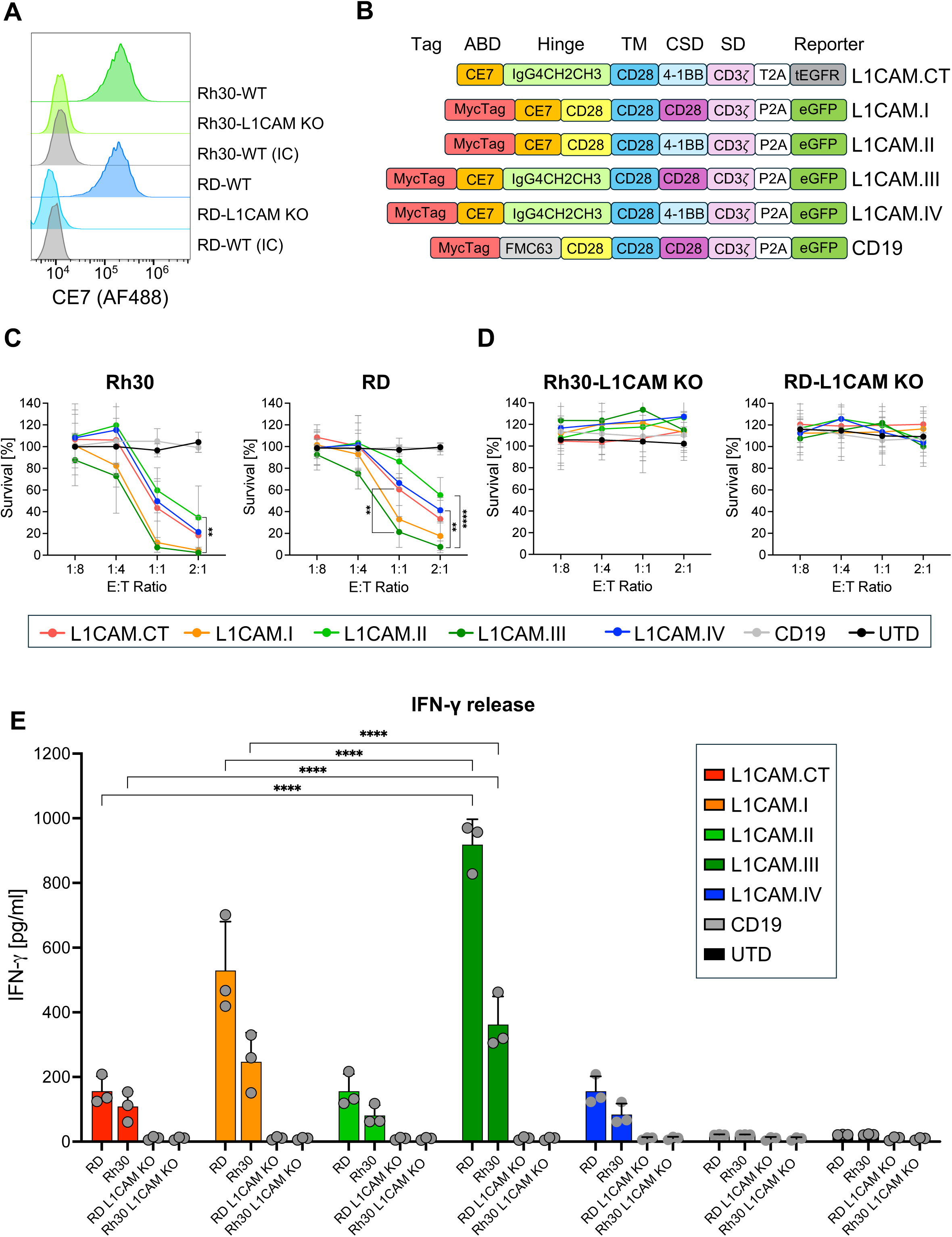
CE7-based CAR constructs selectively recognize L1CAM in RMS. **(A)** Flow cytometry of Rh30 and RD cells (wild-type, L1CAM knockout, and isotype controls) stained with CE7 antibody to detect tumor-associated L1CAM. **(B)** Schematic of CE7-derived CAR constructs with varied hinge and costimulatory domains. Domains shown include antigen-binding domain (ABD), hinge, transmembrane (TM), costimulatory (CSD), signaling domain (SD), and reporter elements. **(C-D)** Tumor cell killing by CAR T cells against fLuc^+^ Rh30 and RD cells **(C)** or their L1CAM knockout counterparts **(D)** after 48 h coculture at E:T ratios of 1:8, 1:4, 1:1, and 2:1, measured by luciferase-based survival assay. Data represent CAR T cells generated from four independent donors (n = 4), each tested in technical triplicates. **(E)** IFN-γ release by CAR T cells cocultured with fLuc^+^ Rh30 and RD cells and corresponding L1CAM knockout targets, measured by ELISA in culture supernatants 24 h post coculture. Data represent CAR T cells generated from three independent donors (n = 3), each tested in technical triplicates. Statistical analysis for panels C-E was performed using two-way ANOVA followed by Dunnett’s multiple comparisons test against L1CAM.III as reference. *P* values are denoted as: ns, not significant; \**p* < 0.05; \*\**p* < 0.01; \*\*\**p* < 0.001; \*\**p* < 0.0001.

Starting from this backbone, we generated four additional CE7-based CAR constructs incorporating different hinge regions or costimulatory domains (Figure 2B). CD19-CAR T cells and untransduced T cells (UTD) served as negative controls. CAR T cells from at least three independent donors were manufactured and characterized (Supplementary Figure S1).^9^

Tumor cell killing was assessed in 48 h coculture assays with Effector to Target (E:T) ratios of 1:8 to 2:1. All L1CAM-CARs induced antigen-specific cytotoxicity in both Rh30 and RD cells, while no effect was observed with CD19-CAR T cells and UTD T cells (Figure 2C). Against Rh30 cells, the long-hinge CD28 CAR (L1CAM.III) provided CAR T cells with the strongest cytotoxicity, reducing tumor survival to <10% at an E:T ratio of 2:1, compared to ∼30% for both L1CAM.CT and L1CAM.II, and ∼20% for L1CAM.IV. A similar pattern was also observed with RD cells, where L1CAM.III-CAR T cells reduced survival to ∼30% at the same ratio, in contrast to ∼60% for L1CAM.CT-, ∼50% for L1CAM.II-, and ∼40% for L1CAM.IV-CAR T cells. At lower E:T ratios, CD28-based constructs (L1CAM.I and L1CAM.III) consistently outperformed 4-1BB-based designs. This difference was observed with both short and long hinge formats, indicating that CD28 costimulation was the dominant factor driving cytotoxicity. CE7-CARs showed no cytotoxicity against L1CAM knockout Rh30 and RD cells, confirming antigen specificity (Figure 2D).

To further assess effector function beyond direct cytotoxicity, we measured IFN-γ secretion 24 h after coincubation of RMS cells with L1CAM-CAR T cells. Among the panel, L1CAM.III-CAR T cells elicited the strongest cytokine release, inducing ∼6-fold and ∼3-fold higher levels than the clinical reference L1CAM.CT-CAR T cells in RD and Rh30 models, respectively (Figure 2E). L1CAM.I-CAR T cells also showed enhanced activity, reaching ∼3.5-fold higher than L1CAM.CT in RD cells, whereas L1CAM.II- and L1CAM.IV-CAR T cells induced only minimal release, comparable to L1CAM.CT. CD19-CAR and UTD T cells showed no detectable secretion. Across constructs, IFN-γ release was consistently higher in RD cells (∼920 pg/mL for L1CAM.III) than in Rh30 cells (∼360 pg/mL for L1CAM.III), while negligible levels were detected when CAR T cells were co-incubated with L1CAM KO target cells.

### L1CAM.III-CAR T cells exhibit enhanced potency and favorable specificity

To benchmark the performance of the optimized L1CAM.III-CAR T cells, we compared their activity to CAR T cells recognizing B7-H3 - a target expressed at high density in pediatric tumors^37^ and known to support CAR T cell-mediated RMS eradication *in vivo*.^9,12^

Flow cytometry analysis confirmed that B7-H3 is present at approximately 5-fold higher levels than L1CAM (∼50’000 versus ∼10’000 molecules per cell) on both RD and Rh30 cell lines, as previously described^9^ (Figure 1B). Despite this marked discrepancy, L1CAM.III-CAR T cells achieved cytotoxicity comparable to B7-H3-CAR T cells across multiple E:T ratios (Figure 3A) and induced significantly greater IFN-γ secretion against RD cells after 24 h coculture (*p* = 0.001) (Figure 3B).

**Figure 3.**
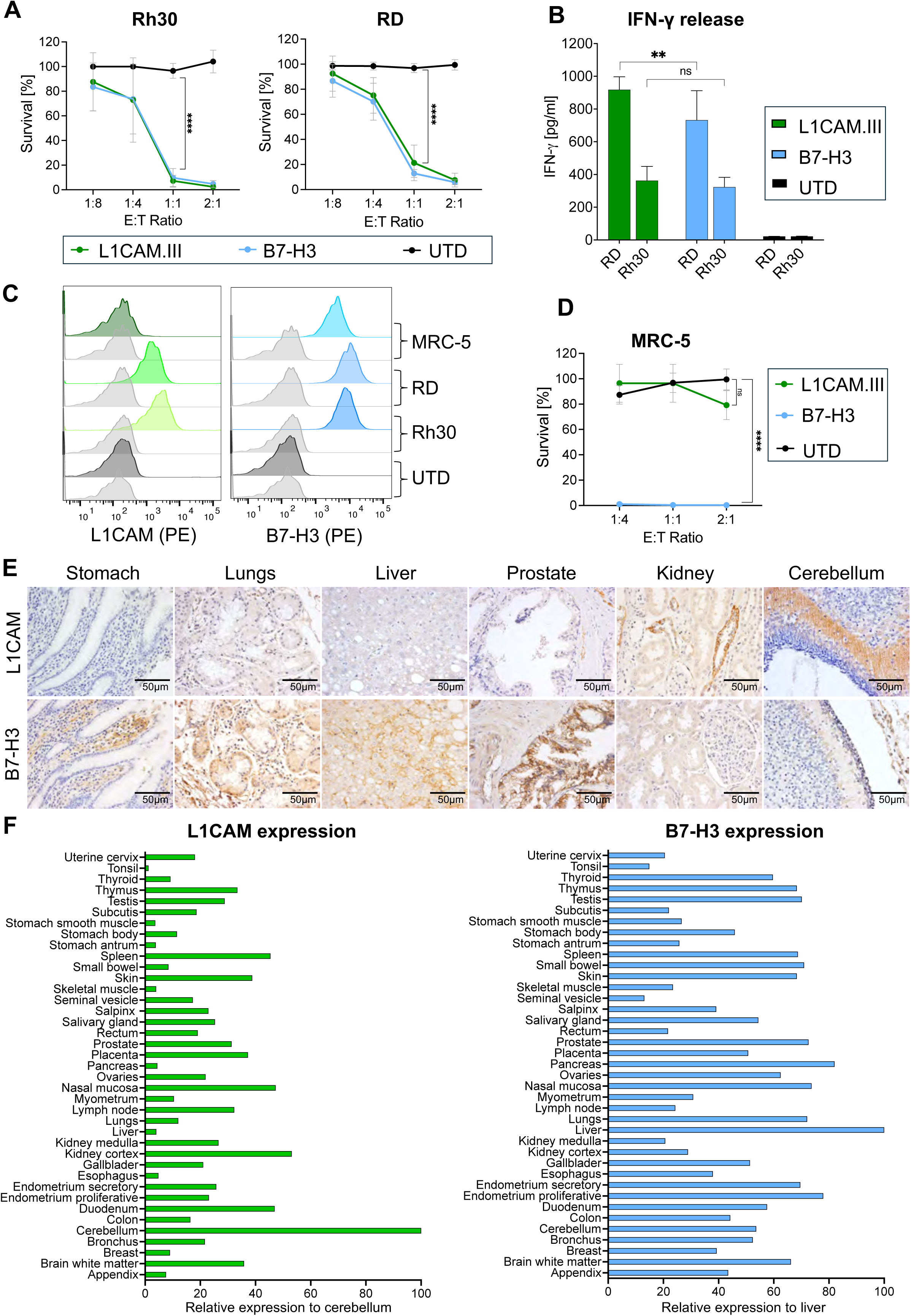
CD28-based L1CAM.III-CAR achieves potency comparable to B7-H3 CARs with improved selectivity. **(A)** Tumor cell killing by L1CAM.III- and B7-H3-CAR T cells against fLuc^+^ Rh30 and RD cells after 48 h coculture at E:T ratios of 1:8, 1:4, 1:1, and 2:1, measured by luciferase-based survival assay. UTD T cells served as negative controls. Data from three independent donors (n = 3), each tested in technical triplicates. **(B)** IFN-γ release from L1CAM.III- and B7-H3-CAR T cells cocultured with Rh30 and RD cells for 24 h, measured by ELISA. Data represent mean ± SD from three independent donors (n = 3), each tested in technical triplicates. **(C)** Flow cytometry showing B7-H3 but not L1CAM expression on MRC-5 fibroblasts. RD and Rh30 cells were used as positive controls, while UTD T cells served as the negative control. Isotype controls are colored in gray. **(D)** Killing of fLuc^+^ MRC-5 fibroblasts by L1CAM.III- and B7-H3-CAR T cells after 48 h coculture, measured by luciferase-based survival assay. Data from three independent donors (n = 3), each tested in technical triplicates. **(E)** IHC of representative healthy human tissues (stomach, lungs, liver, prostate, kidney, cerebellum) stained for L1CAM or B7-H3 (brown) with hematoxylin counterstain (blue). Scale bar = 50 µm. **(F)** Quantitative analysis of IHC staining of TMAs. DAB-positive signal was quantified, with the tissue showing strongest staining set to 100% and all others expressed relative to this maximum. Statistical analysis for panels A, B and D was performed using two-way ANOVA followed by Dunnett’s multiple comparisons test against UTD as reference. *P* values are denoted as: ns, not significant; \**p* < 0.05; \*\**p* < 0.01; \*\*\**p* < 0.001; \*\**p* < 0.0001.

To begin assessing potential off-tumor risks, we tested CAR T cell activity against MRC-5 lung fibroblasts, a widely used primary fibroblast line. Flow cytometry confirmed sustained B7-H3 expression in MRC-5 (Figure 3C), whereas L1CAM was below detection, consistent with our findings (Figure 1). Functionally, B7-H3-CAR T cells mediated potent fibroblast killing even at low E:T ratios, whereas L1CAM.III-CAR T cells showed no activity (Figure 3D), indicating a potentially more favorable safety profile.

To compare the physiological distribution of L1CAM and B7-H3, we stained a panel of healthy tissues and quantified their relative expression. For each antigen, the tissue exhibiting the highest signal was set at 100%, and all other tissues were expressed relative to this reference. L1CAM expression was generally highest in the cerebellum and kidneys, and low and confined to a few compartments in the other tissues tested, consistent with the restricted expression of the CE7 epitope^18,38^ (Figure 3F, left). In contrast, B7-H3 displayed a broader expression pattern, with higher relative expression across multiple organs such as prostate, lung, endometrium, pancreas, and liver, consistent with published results^39^ (Figure 3F, right). Together, these findings show that L1CAM.III-CAR T cells achieve potent antitumor activity *in vitro* while targeting an antigen that is expressed at lower levels than B7-H3 in healthy tissues, indicating higher tumor specificity and potential for safety (Supplementary Figure S2).

### Orthotopic RMS models reveal a superior efficacy of optimized L1CAM.III-CAR T cells over the clinical construct

We next assessed the antitumor activity of L1CAM.CT-, L1CAM.III-, and B7-H3-CAR T cells in orthotopic RMS mouse models. To allow simultaneous *in vivo* monitoring of tumor burden and CAR T cell persistence, RMS cells were engineered to express firefly luciferase (fLuc), while all CAR constructs co-expressed Antares, a NanoLuc-based reporter (Figure 4A). Mice bearing fLuc^+^ Rh30 or RD tumors were treated with two intravenous doses of CAR T cells on days 5 and 12 after tumor inoculation (Figure 4B, G).

**Figure 4.**
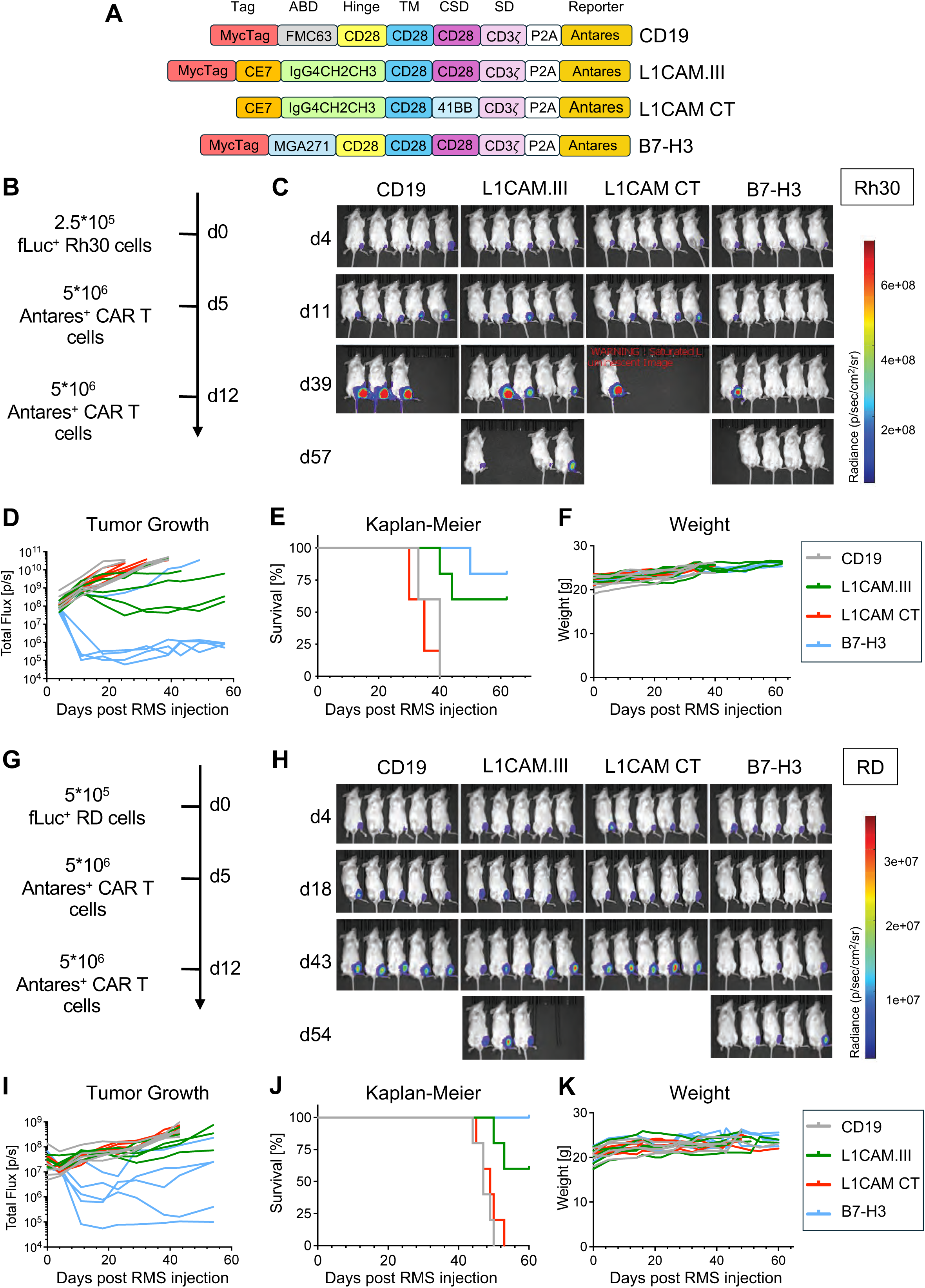
Optimized L1CAM.III-CAR mediates *in vivo* tumor control in orthotopic RMS models. **(A)** Schematic representation of CAR constructs (CD19, L1CAM.III, L1CAM.CT, B7-H3), each co-expressing Antares *via* a P2A linker. **(B)** Experimental setup for Rh30 xenograft study: 2.5×10^5^ fLuc^+^ Rh30 cells injected at day 0, followed by two intravenous doses of 5×10^6^ Antares^+^ CAR T cells on days 5 and 12. **(C)** Representative *in vivo* bioluminescence imaging of Rh30 tumor-bearing mice treated with the indicated CAR T cells (n=5 per group). **(D)** Longitudinal Rh30 tumor growth monitored by bioluminescence flux (photons/s) in individual mice. **(E)** Kaplan-Meier survival analysis of Rh30-bearing mice treated with each CAR T cell group. **(F)** Body weights (g) of Rh30-bearing mice during treatment. **(G)** Experimental setup for RD model: 5×10^5^ fLuc^+^ RD cells injected at day 0, followed by two intravenous doses of 5×10^6^ Antares^+^ CAR T cells on days 5 and 12. **(H)** Representative *in vivo* bioluminescence imaging of RD tumor-bearing mice treated with the indicated CAR T cells (n=5 per group). **(I)** Longitudinal RD tumor growth monitored by bioluminescence flux (photons/s) in individual mice. **(J)** Kaplan-Meier survival analysis of RD-bearing mice treated with the indicated CAR T cells. **(K)** Body weights (g) of RD-bearing mice during treatment.

In the Rh30-derived model, longitudinal bioluminescence imaging revealed no detectable activity of L1CAM.CT-CAR T cells, with tumor progression indistinguishable from CD19-treated controls (Figure 4C-D). In contrast, L1CAM.III-CAR T cells achieved partial response in 3/5 mice with peak bioluminescence reduction of >500-fold at day 25 but not complete remission, while the remaining two mice showed delayed progression (Figure 4C-D). B7-H3-CAR T cells, instead, induced rapid tumor eradication within two weeks of treatment and achieved complete responses in 4/5 mice (Figure 4C-D). Survival was extended in both L1CAM.III- and B7-H3-CAR T cell treated groups compared to CD19- and L1CAM.CT-CAR T cell arms (Figure 4E). Importantly, body weights remained unaltered in all CAR T cell-treated mice (Figure 4F). Likewise, IHC analyses of healthy organs (Supplementary Figure S3) revealed neither unspecific infiltration of CAR T cells nor tissue damage, in line with our finding showing no cross-reaction of CE7 with murine L1CAM (Supplementary Figure S4).

In the RD-derived model, a similar trend in the L1CAM.III- and B7-H3-CAR T cell activity was observed, both outperforming L1CAM.CT- and CD19-CAR T cells (Figure 4H-I). Tumor regression was less pronounced than in Rh30 tumors, with mean bioluminescence reductions of ∼2-fold for L1CAM.III and ∼300-fold for B7-H3 relative to controls. Nevertheless, both L1CAM.III- and B7-H3-CAR T cell groups showed delayed tumor progression compared to CD19- and L1CAM.CT-treated animals, although the magnitude of disease control was modest and ultimately transient in all groups. This partial tumor control translated into a prolonged survival compared with controls (Figure 4J), without weight loss or other signs of toxicity (Figure 4K).

### L1CAM.III- and B7-H3-CAR T cells show enhanced persistence in orthotopic RMS models

Longitudinal bioluminescence imaging of mice bearing Rh30- and RD tumors revealed sustained Antares⁺ CAR T cell signals across multiple anatomical regions throughout the experiment. This finding is consistent with CAR T cells trafficking to lymphoid tissues and tumors. In Rh30-derived model (Figure 5A-E), CAR T cells expanded after each infusion (days 5 and 12) and accumulated in several compartments, with long-term persistence evident beyond 50 days. Region-of-interest analyses demonstrated marked increases in the neck region (Figure 5B) and chest region (Figure 5C) for both L1CAM.III- and B7-H3-CAR T cells, whereas L1CAM.CT remained near baseline. Notably, B7-H3-CAR T cells could not be detected in one Rh30 mouse, which was the only animal in this cohort that failed to respond. Interestingly, CD19-CAR T cells showed detectable signals in the same regions despite their lack of tumor control, which may reflect antigen-independent tonic signaling previously reported for the FMC63-based construct used in our B7-H3 CARs.^40^

**Figure 5.**
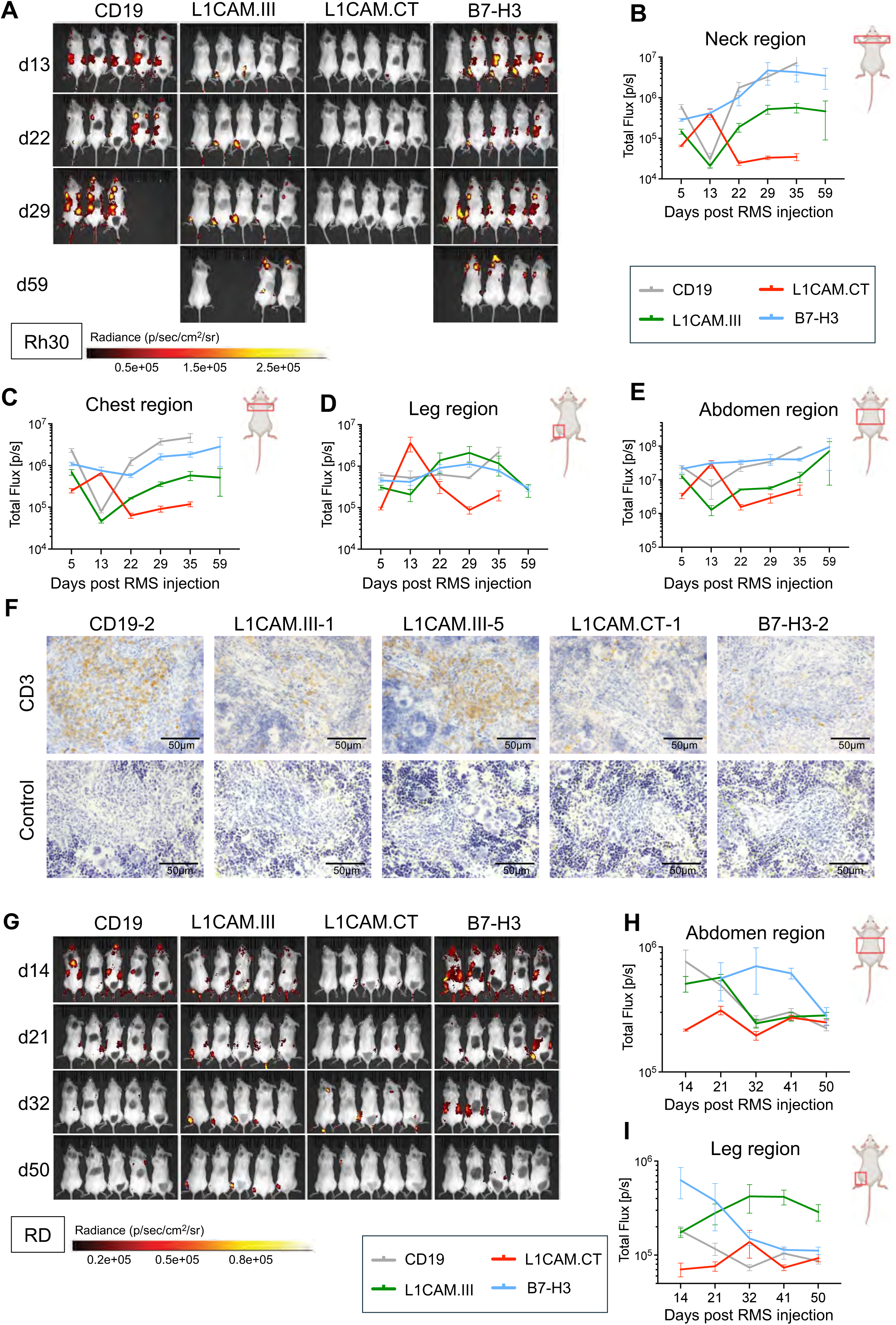
L1CAM.III-CAR T cells persist and traffic to tumor and lymphoid tissues *in vivo.* **(A)** Whole-body bioluminescence imaging of Antares⁺ CAR T cells in Rh30-bearing mice at indicated time points (n=5 per group). **(B-E)** Region-of-interest (ROI) quantification of Antares signal showing CAR T cell expansion in the **(B)** neck (cervical lymph nodes), **(C)** chest (axillary lymph nodes), **(D)** tumor-bearing gastrocnemius muscle, and **(E)** abdominal region. **(F)** IHC of spleens confirming CAR T cell infiltration. Spleens were collected at endpoint, stained for CD3⁺ T cells, showing CAR T cell infiltration. Negative controls (no primary antibody) are included. Staining shows CD3 (brown) and hematoxylin counterstain (blue) Scale bars, 50 µm. **(G)** Whole-body bioluminescence imaging of Antares⁺ CAR T cells in RD-bearing mice at indicated time points (n=5 per group). **(H-I)** ROI quantification of Antares signal in the **(H)** abdominal region and **(I)** tumor-bearing gastrocnemius muscle of RD-bearing mice.

In the leg region where tumors developed (Fig. 5D), L1CAM.CT-CAR T cells homed to the site by day 13 and exhibited a sharp rise in signal, surpassing L1CAM.III- and B7-H3-CAR T cells. This expansion was not sustained, and signal rapidly declined without tumor clearance. In contrast, L1CAM.III-CAR T cells expanded progressively, with kinetics similar to B7-H3-CAR T cells, peaking at day 29 before contracting in association with tumor regression.

In the abdominal region, where the spleen is located (Figure 5E), the strongest signals were detected in mice treated with CD19- or B7-H3-CAR T cells, intermediate levels in those treated with L1CAM.III-CAR T cells, and minimal signals in the L1CAM.CT cohort. IHC of spleens collected at the study endpoint (Figure 5F) indicated that these signals partly reflected CAR T cell homing, as CD3⁺ cells were present in spleens from L1CAM.III- and B7-H3-CAR T cell-treated mice. Staining appeared more intense in animals with higher tumor burden (e.g., L1CAM.III, mouse #5) compared with those achieving near-complete regression (e.g., L1CAM.III, mouse #1; B7-H3, mouse #2). Consistent with the imaging data, CD19, mouse #2 also showed high splenic infiltration, whereas L1CAM.CT-CAR T cell-treated mice exhibited only weak staining.

In the RD-derived model (Figure 5G-I), overall Antares⁺ signals were lower than in Rh30-bearing mice, reflecting reduced CAR T cell persistence. Region-of-interest analyses showed that CD19- and B7-H3-CAR T cells initially exhibited the strongest abdominal signals (likely from the spleen, Figure 5H), whereas L1CAM.III-CAR T cells displayed intermediate levels and L1CAM.CT-CAR T cells remained barely detectable. B7-H3-CAR T cells localized robustly in the tumor-bearing leg (Figure 5I) during early tumor growth but declined as tumors regressed, whereas L1CAM.III-CAR T cells reached moderate intensities and persisted until day 50. In contrast, CD19- and L1CAM.CT-CAR T cells produced only low-level or transient signals in the leg, consistent with poor expansion and lack of tumor control.

## 4. Discussion

RMS remains a major clinical challenge in pediatric oncology, with particularly poor outcomes for patients with FP-RMS and metastatic disease. Despite advances in multimodal therapy, survival for relapsed or metastatic cases remains dismal, highlighting the urgent need for novel treatment strategies.^4^ CAR T cells have revolutionized the treatment of hematologic malignancies,^5,6^ but their application to solid tumors remains limited by the lack of selective antigens and by barriers imposed by antigen density, tumor heterogeneity, and the microenvironment.^41^ In this study, we identify L1CAM as a clinically relevant CAR T cell target in RMS and demonstrate that rational CAR engineering can overcome the challenge of moderate antigen density to achieve potent and selective activity.

By surfaceome profiling, we previously identified L1CAM expression in alveolar RMS.^15^ L1CAM is implicated in tumorigenesis across several cancers,^17^ and displays a restricted expression pattern in healthy tissues, largely confined to neural and renal compartments.^42–44^ In this study, we confirmed consistent L1CAM expression across RMS cell lines and PDX models, with only minimal or focal detection in healthy tissues. Interestingly, while L1CAM expression in patient samples is associate with aRMS,^15^ here we observed substantial expression in eRMS cell lines and PDXs, possibly reflecting selective pressure, epigenetic drift, or engraftment bias in xenografts and long-term cultures. Quantitative profiling revealed that RMS cells express ∼2’000-20’000 L1CAM molecules per cell, placing it in a low-to-intermediate density range compared with B7-H3 (∼50’000/cell) or CD19 (>100’000/cell), emphasizing the need for CAR optimization.

The limited clinical activity of CE7-based CAR T cells observed in the recent neuroblastoma trial (NCT02311621) underscores that the CE7 binder alone might not be sufficient to drive potent clinical response.^23^ Consistent with these findings, our L1CAM.CT construct infiltrated RMS tumors but did not induce objective responses, mirroring the clinical experience. Guided by promising preclinical^18,20^ and clinical^23^ studies of CE7-based CARs, we developed and screened a set of CAR constructs and identified a CE7-scFv coupled to a CD28 costimulatory domain (L1CAM.III) as the most effective design. Consistent with prior work comparing CD28 and 4-1BB signaling domains,^45^ L1CAM.III-CAR mediated the strongest cytotoxicity and IFN-γ secretion compared with alternative CARs, including the one previously tested in the clinical trial (NCT02311621). These results align with prior evidence that CD28 backbones enhance Th1 cytokine production, which can augment efficacy by upregulating MHC and costimulatory molecules in the tumor microenvironment.^46,47^ Notably, L1CAM.III-CAR T cells achieved preclinical activity comparable to B7-H3-directed CAR T cells, despite the lower antigen density of L1CAM. Importantly, *in vitro* studies demonstrated tumor selectivity and absence of off-target effects, highlighting the feasibility of safely targeting a low-expressed antigen.

In contrast, B7-H3-CAR T cells showed a strong cytotoxicity against MRC-5 lung fibroblasts, despite their lower B7-H3 surface density relative to RMS cells. This likely reflects the slow proliferation rate of primary fibroblasts and sensitivity of healthy cell lines, which can increase susceptibility to CAR-mediated killing in short-term assays. These findings emphasize a critical distinction between L1CAM and B7-H3 targeting. While both antigens support potent antitumor activity, L1CAM seems to offer a more favorable selectivity profile with reduced risk of stromal toxicity.

Our comparative *in vivo* studies further confirmed the therapeutic potential of L1CAM.III-CAR T cells, while also revealing RMS subtype-dependent differences. Notably, tumor growth is much faster in the FP-RMS Rh30 model than in the FN-RMS RD model. Interestingly, L1CAM.III-CAR T cells induced partial responses and prolonged survival in the FP-RMS Rh30 model but had more limited activity in the FN-RMS RD model. Similar trends observed with B7-H3-CAR T cells suggest that tumor-intrinsic or microenvironmental mechanisms - such as adenosinergic signaling,^48^ IFN-γ-induced PD-L1 upregulation,^49^ or extracellular matrix remodeling^41^ - may contribute to resistance in FN-RMS, and need to be investigated in the future. A similar response in RD and Rh30 tumors was expected based on their level of L1CAM expression in culture. The observed differences may be due to a different in vivo expression, as is evidenced in the IHC staining of orthotopic xenografts, where L1CAM expression is prominently associated with the cell membrane only in Rh30. L1CAM can be cleaved from the cell surface as a means of resistance and exists in a soluble form known as soluble L1CAM (sL1CAM). This mechanism has been implicated in immune evasion in neuroblastoma and other cancers,^50^ though its role in FN-RMS remains unexplored and merits further investigation. Although L1CAM.III-CAR T cells showed improved *in vivo* expansion, as monitored by nano-luciferase detection, and delayed tumor progression relative to L1CAM.CT in some mice, particularly in the FP-RMS Rh30 model, these improvements were modest and did not prevent eventual relapse. This underscores the need for further optimization to achieve long-lasting responses. The CAR T cells production before infusion could also have an impact on the efficiency downstream, which deserves a particular attention in future studies.

Encouragingly, our candidate construct did not induce overt toxicity in mice, confirming tumor selectivity. These observations are limited by the species differences, as CE7 does not cross-react with murine L1CAM. Indeed, the valuable experience gained in the first in human clinical trial mentioned above underscores the need for caution: while early CE7-based trials reported only mild toxicities, the ENCIT-01 study associated more potent constructs with cytokine release syndrome (CRS) and cutaneous effects.^23^ Although manageable, these adverse effects observed with 4-1BB CSD based constructs, indicate that translation of CD28-based designs will require carefully controlled dose escalation to define the therapeutic window.

Taken together, these findings establish L1CAM as a rational and promising immunotherapy target in RMS, specifically the more aggressive FP-RMS. We demonstrate that rational CAR design can overcome the challenge of moderate antigen density, achieving potent activity while preserving selectivity for a solid tumor. This work provides a strong foundation for further development of L1CAM-directed CAR T cells in RMS based on a logic-gating approach, while underscoring the need to address tumor heterogeneity, microenvironmental barriers, and safety in future studies. Ultimately, these results open a new avenue for targeting solid tumors that could expand the therapeutic arsenal against pediatric cancers.

## Declarations

### Authors’ contributions

Conceptualization: CP, AT, MB. Funding acquisition: CP, RF, JR, MB. Investigation: CP, SGD, CG, AT. Methodology: CP, CG, AT, MB. Supervision: JR, MB. Writing - original draft: CP. Writing - review & editing: CP, CG, SGD, RF, JR, AT, MB.

### Ethics approval

The mice experiments were approved by Animal Welfare Office of Canton Bern and in accordance with Swiss Federal Law (Authorization BE22/2022).

### Data availability statement

The data used in this study are available from the lead contact upon request.

### Declaration of generative AI use

During the preparation of this work the author(s) used ChatGPT in order to improve the text. After using this tool, the authors reviewed and edited the content as needed and take full responsibility for the content of the published article.

## Acknowledgments

We are also very grateful to Dr. Stephan Müller and the Flow Cytometry and Cell Sorting (FCCS) facility of the DBMR (University of Bern) for their support. We kindly acknowledge Dr. Carlotta Detotto and the Central Animal Facility (CAF) of the University of Bern for their support. We are grateful to Prof. Andreina Schöberlein and her lab for their help with immunohistochemistry. We also thank Prof. Michael C. Jensen (Seattle Children’s Research Institute) for generously providing the L1CAM CAR plasmid, and Prof. Annette Künkele (Charité Berlin) for helpful discussions and support.

---

**Supplementary Figure S1.**
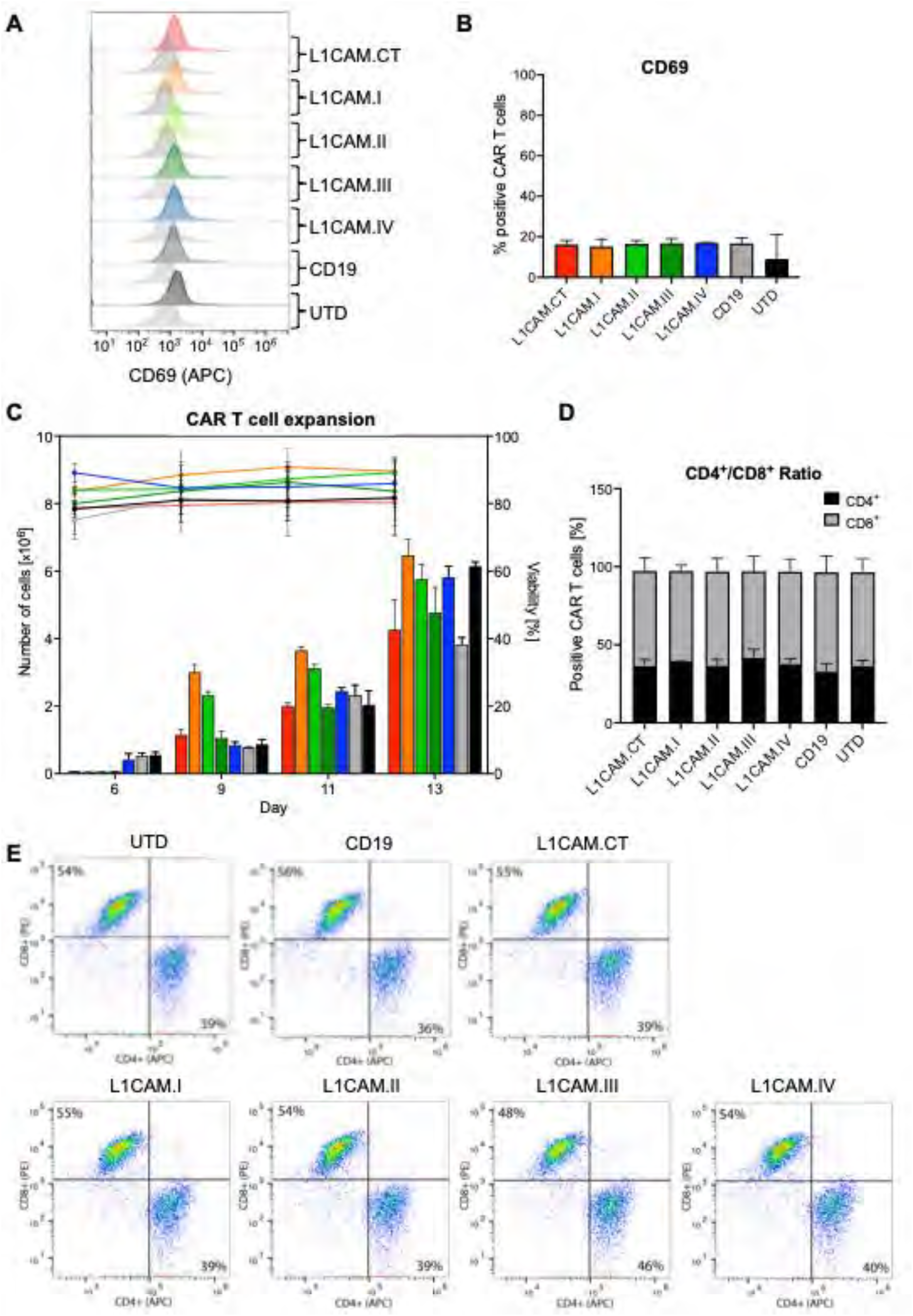
Characterization and expansion of L1CAM-CAR T cells. **(A)** Flow cytometry analysis of CD69 expression on CAR T cells transduced with L1CAM.CT, L1CAM.I, L1CAM.II, L1CAM.III, L1CAM.IV, CD19 CAR, or left untransduced (UTD). Representative histograms are shown. **(B)** Quantification of CD69 expression (% positive CAR T cells) across constructs. Data represent mean ± SD from three independent donors (n=3). **(C)** CAR T cell expansion over 13 days in culture. Cell counts (left y-axis, bars) and viability (right y-axis, lines) are shown. Data represent mean ± SD from three independent donors (n=3). **(D)** CD4/CD8 ratio of CAR T cells determined by flow cytometry on day 14. Data represent mean ± SD from three independent donors (n=3). **(E)** Representative flow cytometry plots showing CD4 and CD8 distribution among UTD, CD19, and L1CAM CAR T cells (CT, I, II, III, IV) on day 14.

**Supplementary Figure S2.**
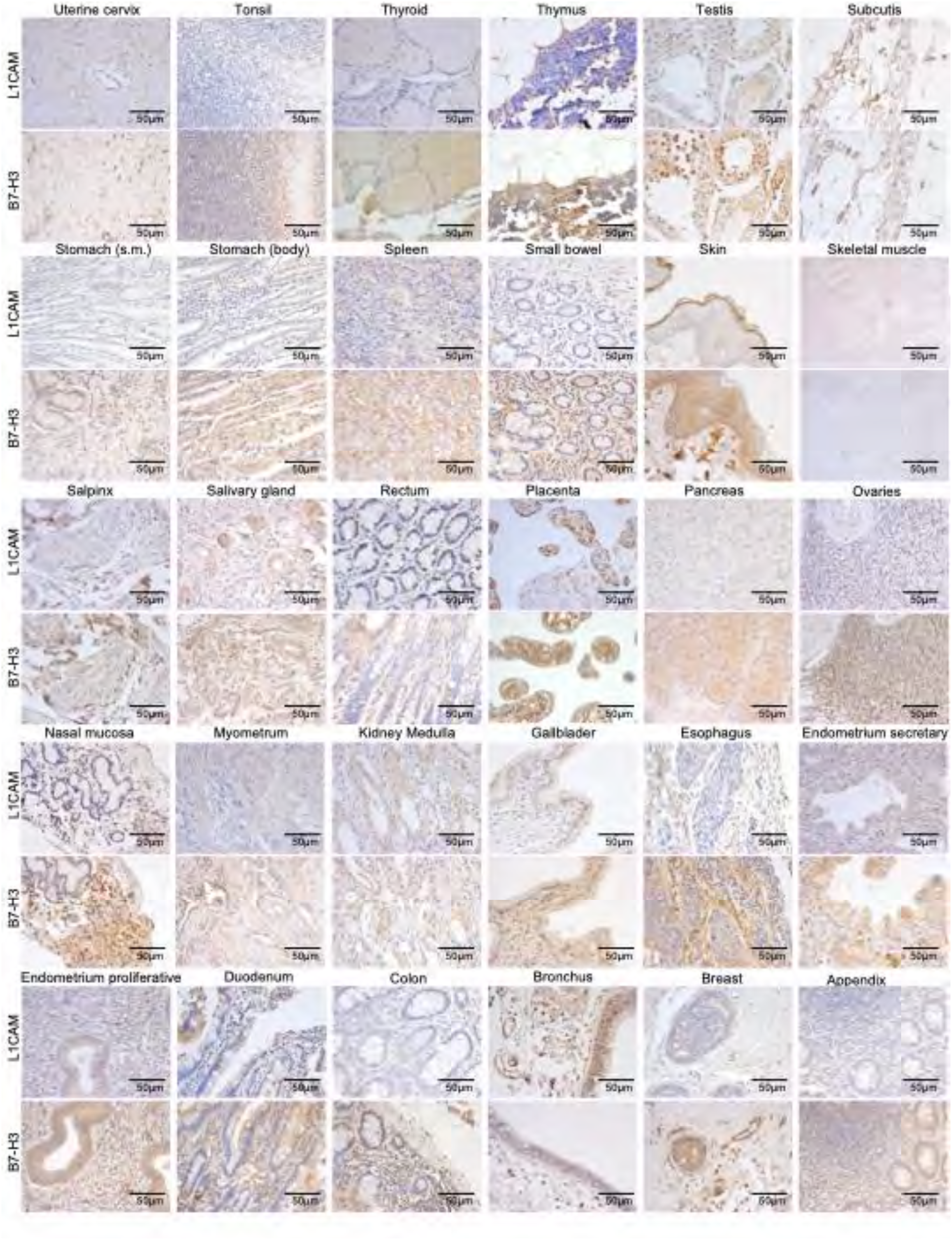
L1CAM and B7-H3 expression in normal tissues. Immunohistochemical staining of L1CAM and B7-H3 across a panel of normal human tissues. Formalin-fixed paraffin-embedded sections were stained with anti-L1CAM antibody (upper rows) or B7-H3 antibody (lower rows) and counterstained with hematoxylin. Scale bars: 50 μm.

**Supplementary Figure S3.**
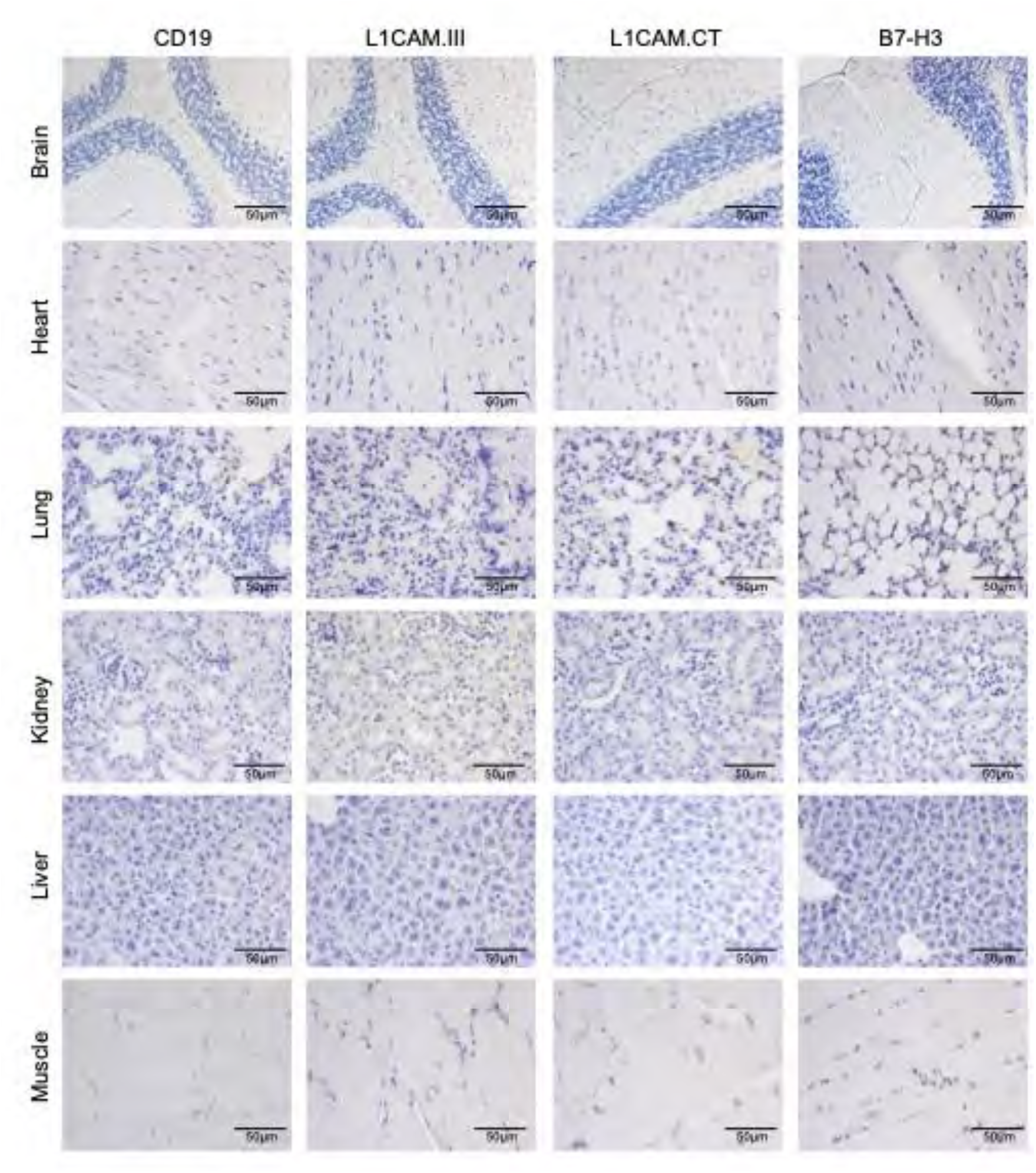
Hematoxylin staining of mouse organs after CAR T cell treatment. Formalin-fixed paraffin-embedded sections were prepared from brain, heart, lung, kidney, liver, and skeletal muscle of mice treated with CD19-, L1CAM.III-, L1CAM.CT-, or B7-H3-CAR T cells. Sections were stained with hematoxylin to assess general tissue morphology and cellular architecture. Across all treatment groups, no histological abnormalities, necrosis, or inflammatory infiltrates were observed. Representative images are shown for each organ and CAR T cell condition. Scale bars: 50 µm.

**Supplementary Figure S4.**
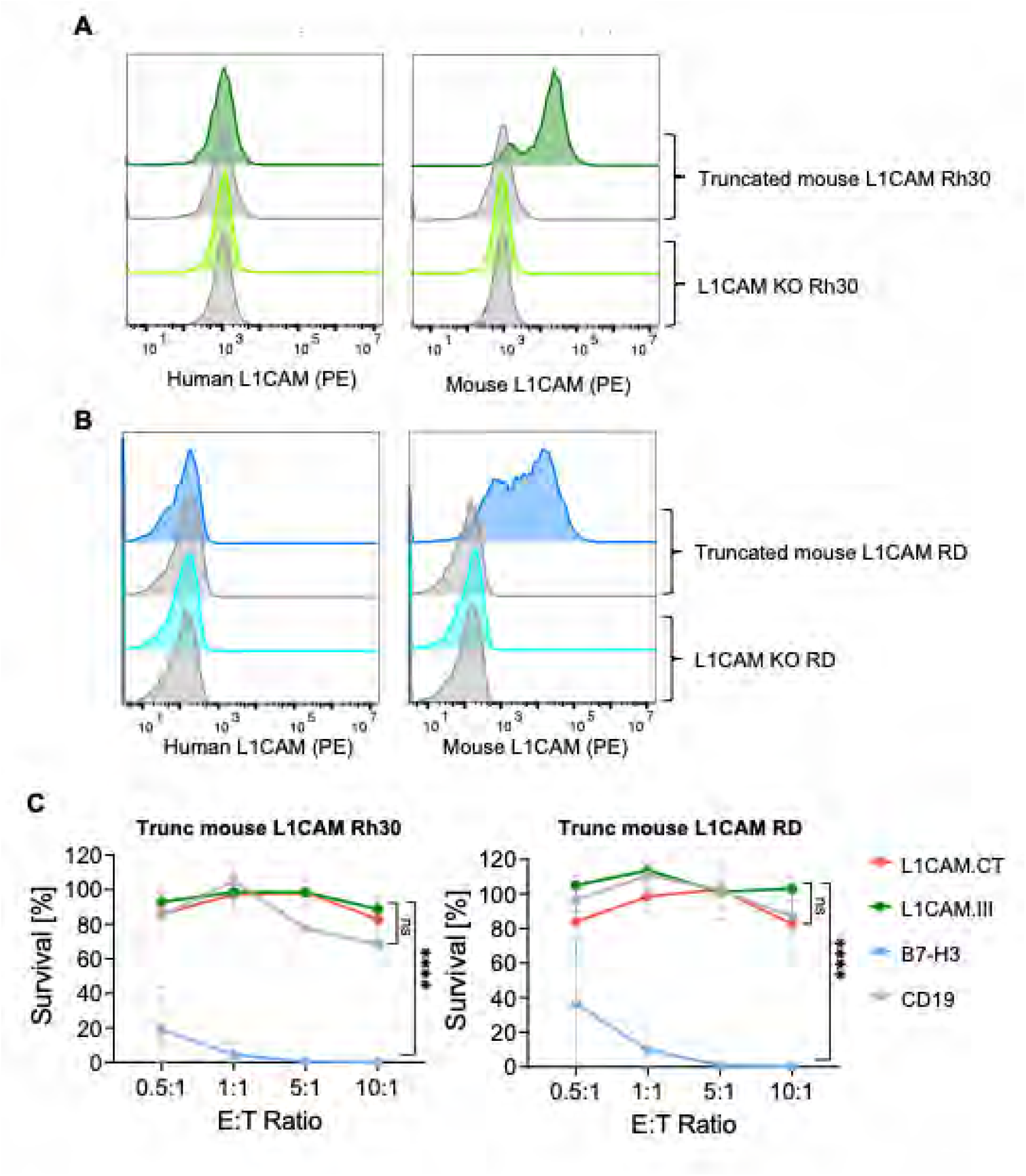
Antigen specificity and functionality of L1CAM CAR T cells. **(A-B)** Flow cytometry analysis of human and mouse L1CAM expression in Rh30 **(A)** and RD **(B)** cells. L1CAM knockout (KO) cells were transduced to express truncated mouse L1CAM. Gray histograms represent isotype controls. **(C)** Tumor cell killing by L1CAM.III- and B7-H3-CAR T cells against fLuc⁺ Rh30 (left) and RD (right) cells after 48 h co-culture at E:T ratios of 1:2, 1:1, 5:1, and 10:1, measured by luciferase-based survival assay. CD19-CAR T cells served as negative controls. Data represent CAR T cells from three independent donors (n=3), each tested in technical triplicates. Statistical analysis for panels A, B and D was performed using two-way ANOVA followed by Dunnett’s multiple comparisons test against CD19-CAR T cells as reference. P values are denoted as: ns, not significant; \**p* < 0.05; \*\**p* < 0.01; \*\*\**p* < 0.001; \*\**p* < 0.0001.

