## Supplementary Figures for "L1CAM-CAR T cells with enhanced potency overcome low-density antigen expression in rhabdomyosarcoma"

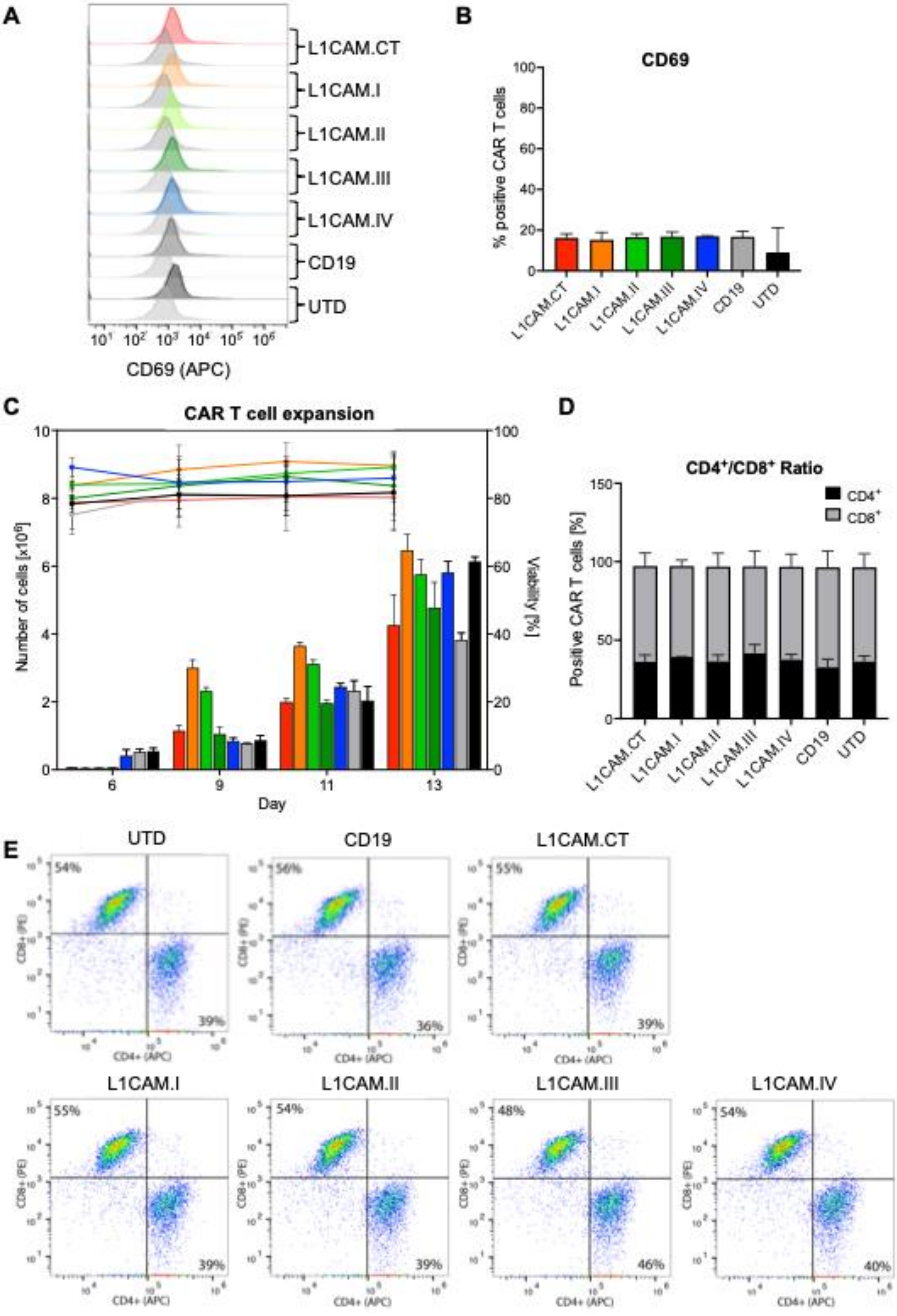

A
L1CAM.CT
L1CAM.I
L1CAM.II
L1CAM.III
L1CAM.IV
CD19
UTD
CD69 (APC)
B
CD69
% positive CAR T cells
L1CAM.CT
L1CAM.I
L1CAM.II
L1CAM.III
L1CAM.IV
CD19
UTD
C
CAR T cell expansion
Number of cells [x10^6]
Viability [%]
Day
6
9
11
13
D
CD4+/CD8+ Ratio
CD4+
CD8+
Positive CAR T cells [%]
L1CAM.CT
L1CAM.I
L1CAM.II
L1CAM.III
L1CAM.IV
CD19
UTD
E
UTD
54%
39%
CD19
56%
36%
L1CAM.CT
55%
39%
L1CAM.I
55%
39%
L1CAM.II
54%
39%
L1CAM.III
48%
46%
L1CAM.IV
54%
40%
CD8+ (PE)
CD4+ (APC)

**Supplementary Figure S1. Characterization and expansion of L1CAM-CAR T cells.** **(A)** Flow cytometry analysis of CD69 expression on CAR T cells transduced with L1CAM.CT, L1CAM.I, L1CAM.II, L1CAM.III, L1CAM.IV, CD19 CAR, or left untransduced (UTD). Representative histograms are shown. **(B)** Quantification of CD69 expression (% positive CAR T cells) across constructs. Data represent mean ± SD from three independent donors (n=3). **(C)** CAR T cell expansion over 13 days in culture. Cell counts (left y-axis, bars) and viability (right y-axis, lines) are shown. Data represent mean ± SD from three independent donors (n=3). **(D)** CD4/CD8 ratio of CAR T cells determined by flow cytometry on day 14. Data represent mean ± SD from three independent donors (n=3). **(E)** Representative flow cytometry plots showing CD4 and CD8 distribution among UTD, CD19, and L1CAM CAR T cells (CT, I, II, III, IV) on day 14.

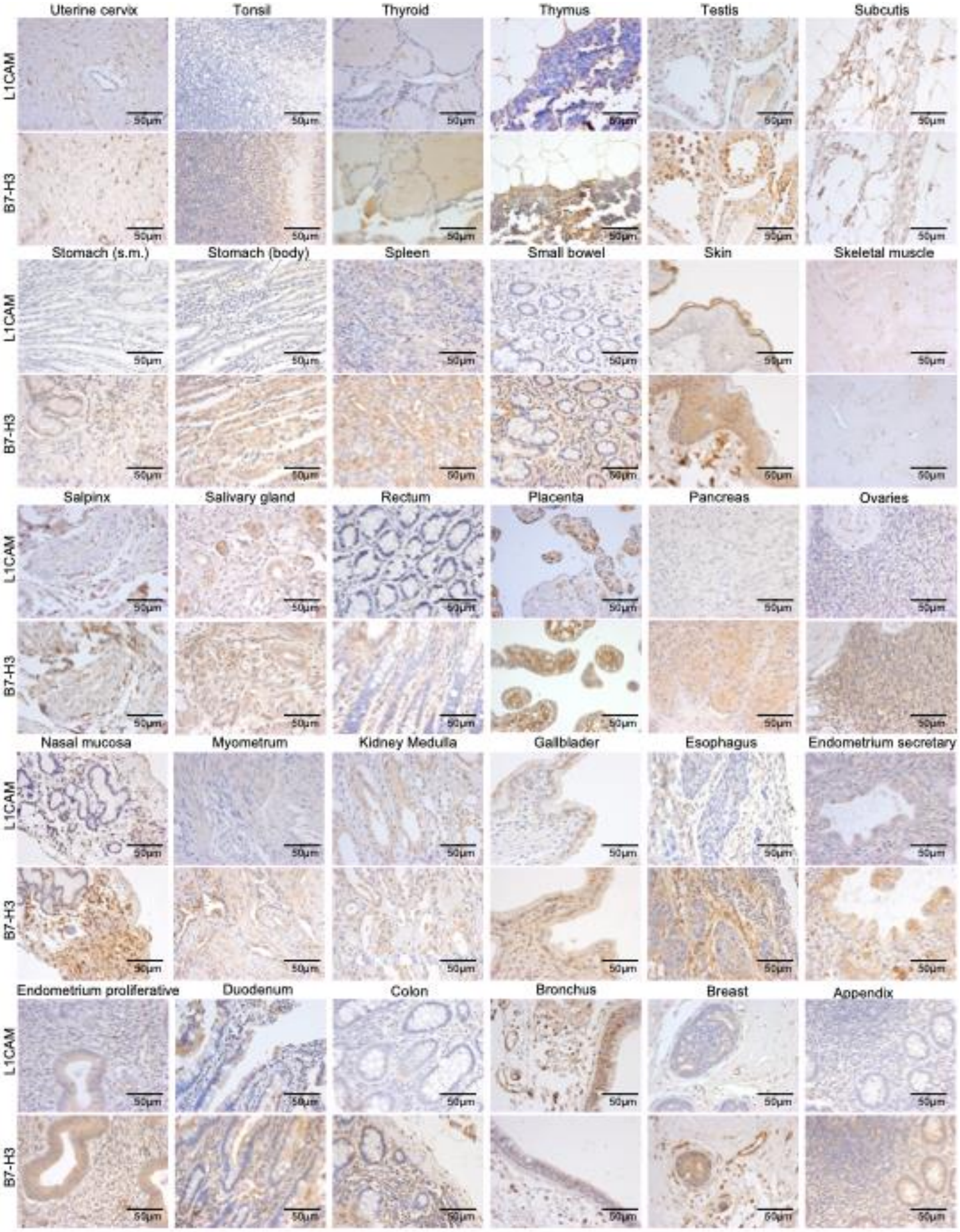

Uterine cervix
Tonsil
Thyroid
Thymus
Testis
Subcutis
L1CAM
B7-H3
Stomach (s.m.)
Stomach (body)
Spleen
Small bowel
Skin
Skeletal muscle
Salpinx
Salivary gland
Rectum
Placenta
Pancreas
Ovaries
Nasal mucosa
Myometrum
Kidney Medulla
Gallblader
Esophagus
Endometrium secretary
Endometrium proliferative
Duodenum
Colon
Bronchus
Breast
Appendix
50μm

**Supplementary Figure S2. L1CAM and B7-H3 expression in normal tissues.** Immunohistochemical staining of L1CAM and B7-H3 across a panel of normal human tissues. Formalin-fixed paraffin-embedded sections were stained with anti-L1CAM antibody (upper rows) or B7-H3 antibody (lower rows) and counterstained with hematoxylin. Scale bars: 50 μm.

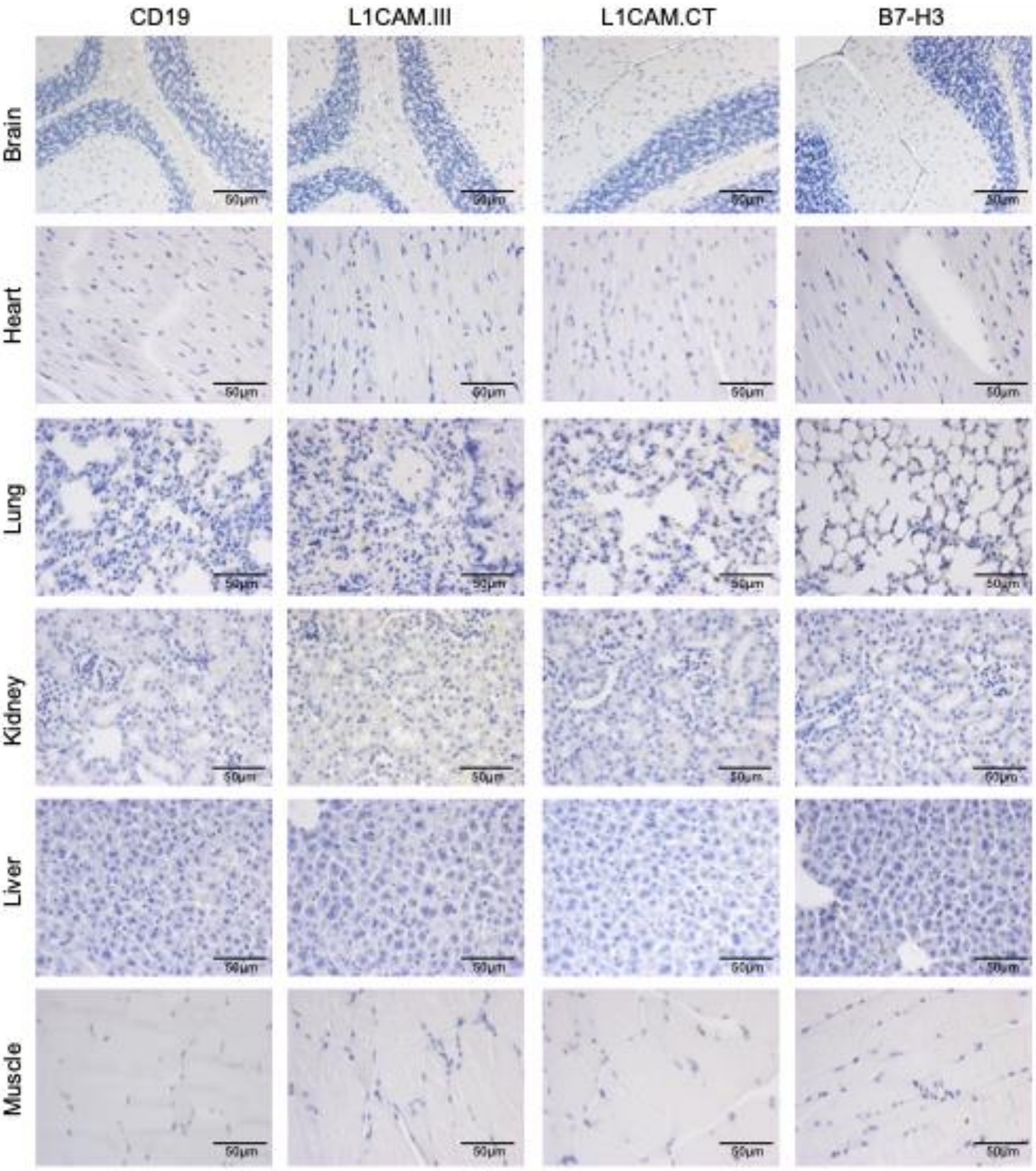
CD19
L1CAM.III
L1CAM.CT
B7-H3
Brain
Heart
Lung
Kidney
Liver
Muscle
50μm

**Supplementary Figure S3. Hematoxylin staining of mouse organs after CAR T cell treatment.** Formalin-fixed paraffin-embedded sections were prepared from brain, heart, lung, kidney, liver, and skeletal muscle of mice treated with CD19-, L1CAM.III-, L1CAM.CT-, or B7-H3-CAR T cells. Sections were stained with hematoxylin to assess general tissue morphology and cellular architecture. Across all treatment groups, no histological abnormalities, necrosis, or inflammatory infiltrates were observed. Representative images are shown for each organ and CAR T cell condition. Scale bars: 50 μm.

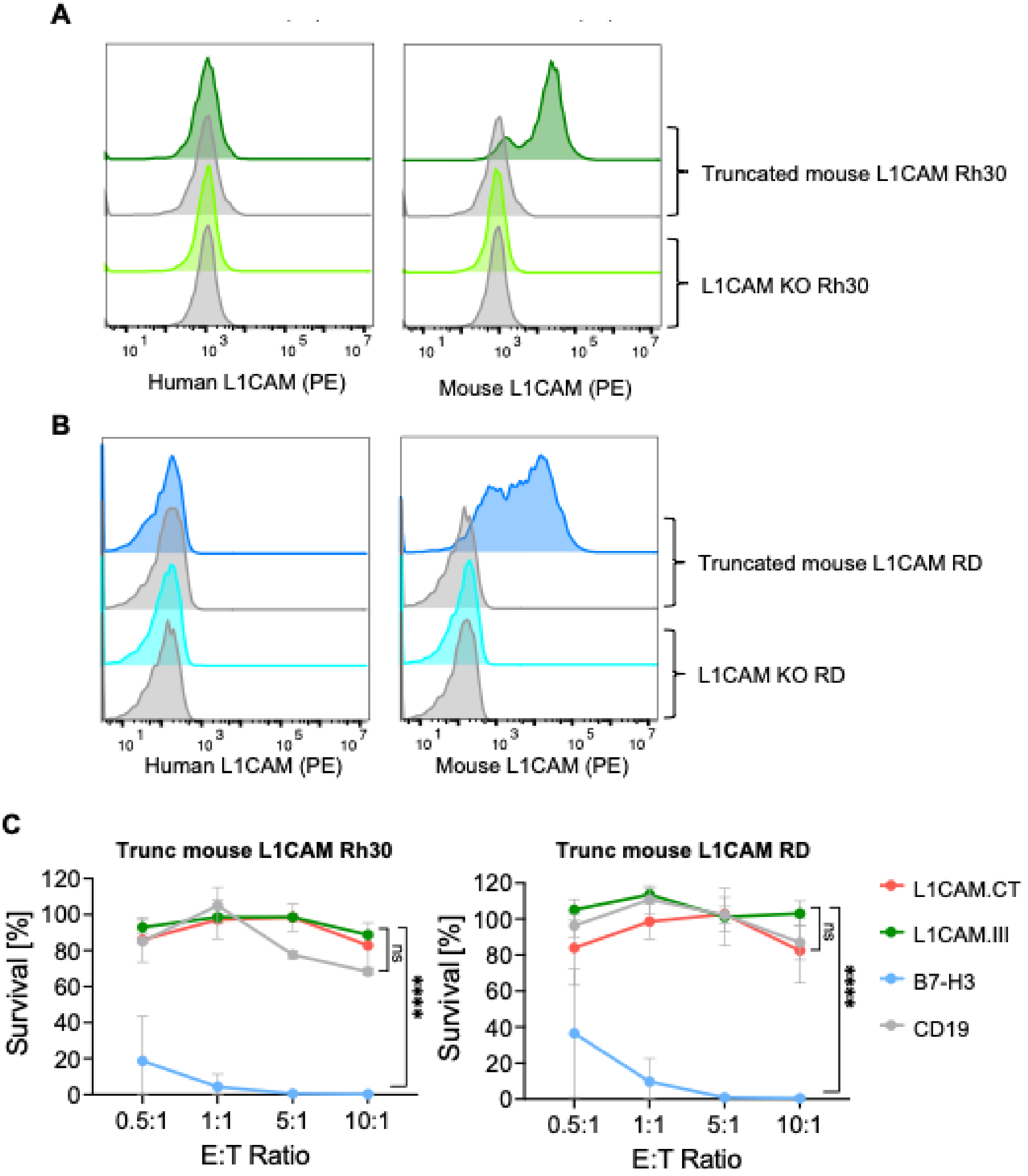
A
Truncated mouse L1CAM Rh30
L1CAM KO Rh30
Human L1CAM (PE)
Mouse L1CAM (PE)
B
Truncated mouse L1CAM RD
L1CAM KO RD
Human L1CAM (PE)
Mouse L1CAM (PE)
C
Trunc mouse L1CAM Rh30
Trunc mouse L1CAM RD
Survival [%]
E:T Ratio
0.5:1
1:1
5:1
10:1
ns
****
L1CAM.CT
L1CAM.III
B7-H3
CD19

**Supplementary Figure S4. Antigen specificity and functionality of L1CAM CAR T cells.** **(A-B)** Flow cytometry analysis of human and mouse L1CAM expression in Rh30 **(A)** and RD **(B)** cells. L1CAM knockout (KO) cells were transduced to express truncated mouse L1CAM. Gray histograms represent isotype controls. **(C)** Tumor cell killing by L1CAM.III- and B7-H3-CAR T cells against fLuc$^{+}$ Rh30 (left) and RD (right) cells after 48 h co-culture at E:T ratios of 1:2, 1:1, 5:1, and 10:1, measured by luciferase-based survival assay. CD19-CAR T cells served as negative controls. Data represent CAR T cells from three independent donors (n=3), each tested in technical triplicates. Statistical analysis for panels A, B and D was performed using two-way ANOVA followed by Dunnett's multiple comparisons test against CD19-CAR T cells as reference. P values are denoted as: ns, not significant; *$p < 0.05$; **$p < 0.01$; ***$p < 0.001$; **$p < 0.0001$.
